# Beta bursts correlate with synchronization of movements to rhythmic sounds

**DOI:** 10.1101/2023.03.14.532353

**Authors:** Qiaoyu Chen, Craig J. McAllister, Mark T. Elliott, Kimron L. Shapiro, Simon Hanslmayr

**Author notes:** **Contact Information / Corresponding Author**, Simon Hanslmayr.

## Abstract

Accumulating evidence indicates transient beta bursts play an important role in the representation of temporal information and prediction. However, the role of beta bursts in sensorimotor synchronization (SMS) involving active interactions between motor and sensory systems to synchronize predictive movements to periodic events remains unclear. To answer this question, 15 participants were invited to complete a finger-tapping task whilst high-density EEG (128 channels) was recorded. Participants tapped with their right index finger in synchrony with 1 Hz and 0.5 Hz tone trains. In line with previous findings, we found a negative mean asynchrony between tone and tap time, i.e., taps preceded tones for both tone frequencies (1 and 0.5 Hz). In the EEG data, beta bursts were detected and their timing in relationship with tapping and auditory tracking was examined. Results revealed that beta bursts tracked tapping and were modulated by the low frequency phase of the tone frequency (i.e., 1 Hz or 0.5 Hz). Importantly, the locking of beta bursts to the phase of auditory tracking correlated with the behavioural variance on a single trial level that occurred while tapping to the tones. These results demonstrate a critical role for an interplay between beta bursts and low frequency phase in coordinating rhythmic behaviour.

## 1. Introduction

Sensorimotor synchronization (SMS), defined as the coordination of movement to external periodic events (Pressing, 1999; Repp, 2005; Repp & Su, 2013), is argued to play an important role in the evolution of music and even of language (Merker, 2000; Colling et al., 2017; Fujii & Wan, 2014; Park et al., 2016). Predictable rhythms optimize rhythmic movement. This improvement in behaviour cannot be explained by a reduction in reaction time but instead is a reflection of predictive timing, as the time gap between movement and auditory stimuli is shorter than the fastest reaction time (Repp, 2005; Repp & Su, 2013). Finger-tapping to an isochronous auditory metronome is amongst the most popular paradigms to investigate SMS because of their simplicity. These tasks usually show that the tap occurs before the tone, often referred to as negative mean asynchrony (NMA; Repp, 2005; Repp & Su, 2013). Moreover, synchronising movements to auditory cues has been applied in motor therapies for movement disorders caused by stroke and cerebral palsy (Thaut et al., 1997; Wright et al., 2014; Wright et al., 2017). However, how the sensory and motor systems interact, and support SMS with temporal predictions, remains uncertain.

Sensory entrainment occurs when ongoing brain oscillations in the stimulated region adapt to the phase of sensory rhythmic stimuli (Hanslmayr et al., 2019; Lakatos et al., 2008, 2019; Wang et al., 2018; Chen et al., 2021). Sensory entrainment has been proposed to coordinate temporal predictions by temporally modulating the excitability of relevant neural assemblies, thus optimizing perceptual analysis of periodic stimuli (Schroeder & Lakatos, 2009; Large & Jones, 1999; Stefanics et al., 2010; Cravo et al., 2011; but see Breska & Deouell, 2017).

Besides its role in temporal prediction, entrainment was observed to facilitate movement (but see Morillon et al., 2016 and Cope et al., 2012 for dissociating the role of periodic stimuli in temporal prediction), especially in the auditory domain (Arnal, 2012; Arnal & Giraud, 2012; Lakatos et al., 2019; Schroeder & Lakatos, 2009). Auditory entrainment may improve movements by affecting the motor system. Passive listening studies confirmed this assertion by observing that the motor system, which serves as a core network that supports timing and time perception, is involved in beat and rhythm processing (Fujioka et al., 2012; Grahn & Brett, 2007; Morillon & Baillet, 2017). Evidence from studies involving rhythmic movements, however, suggests an opposite direction of influence, where overt movements enhance the perception and precise temporal anticipation of upcoming auditory stimuli (Morillon et al., 2014; Morillon & Baillet, 2017). Critically, SMS has been shown to be associated with bursts of beta oscillations (18-24 Hz) directed to the auditory system from the left sensorimotor cortex in a functional connectivity study (Morillon & Baillet, 2017). Together with the enhancement from entrainment, these findings collectively suggest that the auditory and motor systems actively interact during auditory-cued tasks involving SMS.

Although the timing of beta bursts correlates with reaction time (Little et al., 2019), it is unclear whether it plays a role in the coordination between auditory and motor systems during SMS. To answer this question, we tested whether the timing of beta bursts phase-locked to the auditory stimulation frequency reflects functional coordination between auditory and motor systems in SMS. The relationship between beta bursts and movement timing and auditory tracking was examined in an EEG study while participants performed a finger tapping task. The term entrainment has recently been hotly debated because it assumes the interaction between a physiologically meaningful brain rhythm with an externally presented rhythm which is difficult to establish (Capilla et al., 2011; Hanslmayr et al., 2019; Helfrich et al., 2019). Therefore, we here prefer the term tracking over entrainment to simply denote the locking of brain activity to an external auditory rhythm, thus being agnostic about the underlying neural mechanisms that give rise to that phase locking.

## 2. Materials and Method

### 2.1 Participants

Fifteen healthy participants were recruited (mean age = 21.07; age range = 18-32; five males; all right-handed) and completed the experiment, whereas three were excluded because of a failure to be able to define the frequency of interest for beta burst detection (see below). This left twelve participants reported in all data analysis (mean age = 21.08; age range = 18-32; three males). All participants were compensated with £8 per hour. Ethical approval (ERN14-0950) was obtained from the Research Ethics Committee at the University of Birmingham.

### 2.2 Experimental design and procedure

Metronome beats were presented isochronously in two frequency conditions. One contained 60 tones per trial at 1 Hz and the number of tones was 30 in the other condition at 0.5 Hz. These two conditions corresponded to an interstimulus interval (ISI) of 1 and 2 seconds, respectively (Fig. 1). Hence, they are designated as ISI1 and ISI2 in descriptions of later sessions for convenience. For the same purpose, trials locked to each tap/tone were assigned as *tap/tone trials*, while trials locked to the first tone in a stimulus train were referred to as *trials*.

**Fig. 1.**
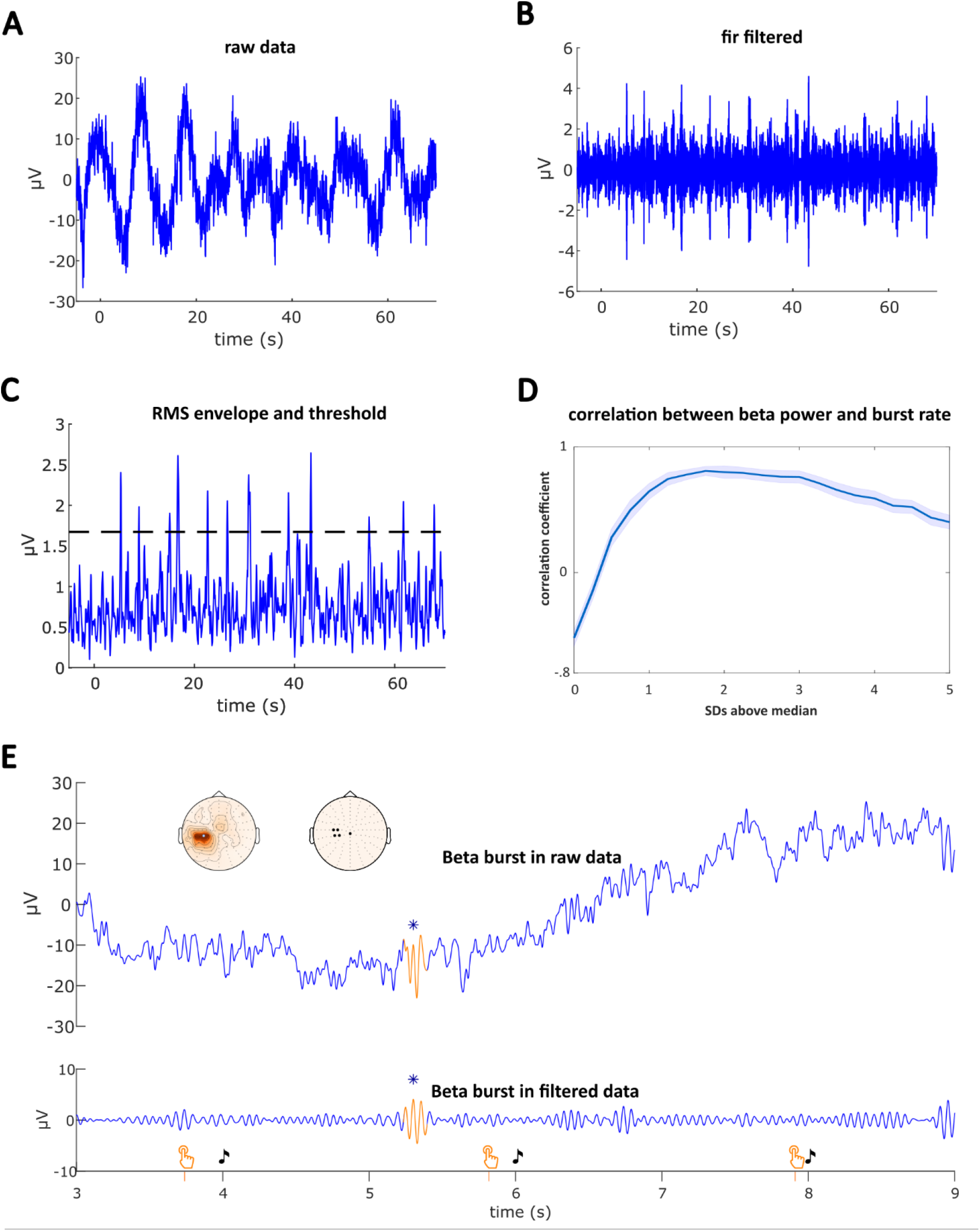
Procedure of algorithm Detection. **(A)** One trial of raw data locked at the first tone of a train. **(B)** Bandpass filtered of the trial with band width of ± 3 Hz centred at individual frequency of interest (18.64 Hz in this example). **(C)** The root mean square (RMS) of the filtered signal. Activity remains above the threshold (the dash line) for 0.1 to 0.5 second were detected as beta burst events. **(D)** The correlation between burst rate and beta power peaked at 1.75 SD above the median of the RMS envelope. **(E)** A detected beta burst event (orange) showed in a zoomed in window from three to nine seconds in the same trial. The orange finger tapping signs denote tap time, while the music notes sign tone onset time. The maximum peak of a detected beta burst was marked by star, both for raw data (upper panel) and filtered data (lower panel). The left topography illustrated the curve fitting resulted ROI for this example, and the right topography demonstrated the ROIs for all participants. Overlapping ROIs were represented by a single dot; one ROI was in electrode A2, two ROIs were in D18, three ROIs were in D17, D19, D28, respectively (128 channel Biosemi layout).

The experiment comprised 60 trials, with 30 of ISI1 and 30 of ISI2. To ensure participants were able to maintain attention, the 60 trials were presented equally in six blocks, with each containing five trials of ISI1 and five trials of ISI2, with a short rest in between blocks. Trials within each block were randomized. Each trial started with a 15s rest period which was used as a baseline for EEG analysis.

Before the experiment, participants were seated in an EEG testing room and instructed to complete an EEG screening form. After the acquisition of consent, preparation of EEG capping and task briefing, participants were seated in a comfortable position with their right hand in a pronated position resting on a wooden pad incorporating a force sensor. Auditory stimuli were delivered through a regular closed back headphone (Sennheiser, Germany). Participants were instructed to tap onto the force sensor with their right index finger in synchrony with the tones. Finger taps and timing of tones were recorded using a National Instruments data acquisition card (DAQ) controlled by the MatTAP toolbox (Elliott et al., 2009) in a MATLAB environment (The MathWorks, Inc., Natick, MA, USA). At the end of experiments, 3D geometric locations of each electrode were recorded by a Polhemus FASTRAK device (Colchester, Vermont, USA) and Brainstorm (Tadel_2011) in MATLAB.

### 2.3 Data analysis

#### 2.3.1 Behavioural Analysis

Descriptive statistics, including mean and standard deviation of the asynchrony between each tap and corresponding tone time (tone minus tap time) were calculated by in-house scripts written in MATLAB (R2018a; The MathWorks, Inc., Natick, MA, USA). This analysis was implemented for ISI1 and ISI2 separately. To evaluate the evolution of adaptation during trials, the asynchrony was first averaged across tap/tone trials at the same time position of a trial for each participant. The group mean was calculated by averaging resulting asynchrony dynamics across participants.

#### 2.3.2 EEG Acquisition and Preprocessing

EEG data was recorded with a 128-channel BioSemi ActiveTwo system. Three extra electrodes were placed at 1 centimetre to the left and right from the lateral canthus, and 1 centimetre below the left eye. Online data were sampled at 2048 Hz via the BioSemi ActiView software.

The preprocessing and analysis of EEG data was conducted in the Fieldtrip toolbox (Oostenveld et al., 2011) in MATLAB. EEG data was first epoched to the first tone of each trial, with 5 s before and 70 s after the onset. The epoched trials were then filtered with a low pass filter of 40 Hz, and a high pass filter of 0.1 Hz. The sampling rate was reduced to 500 Hz before ICA (independent component analysis), followed by manual rejection of noisy channels and trials. ICA components indicating cardiac activity and eye movements were removed. Next, channel interpolation implemented triangulation of the nearest neighbours by calculating the individually recorded electrode locations. Finally, data were re-referenced to the average, and trials with artefacts were rejected by manual inspection.

#### 2.3.3 Beta Burst Detection

##### Defining parameters for burst detection

Before applying the algorithm for beta burst detection, the frequency of interest (FOI) and channel of interest were defined for each participant. To this end, EEG data locked to the tap after pre-processing was first resampled to 250Hz, in the time interval 1 s before and after the tap onset. Then, baseline corrected (‘relchange’, relative change to mean power) power was calculated using wavelet time frequency transformation based on convolution in the time domain, from 1 to 30 Hz in steps of 0.5 Hz with a time resolution of 50 ms at each channel. The FOI for beta burst detection was determined as the frequency in the beta band that was primarily modulated by the tone/tapping rate. Hence, the resulting baseline corrected power data from the previous step was fit to a sine wave at the auditory stimuli rate using the *fit* function in the Curve Fitting Toolbox in MATLAB (R2018a; The MathWorks, Inc., Natick, MA, USA). Specifically, fitting was performed from 10 to 30 Hz (from 15-30 Hz for two participants because of a strong alpha peak at 10-11 Hz) in steps of 0.5 Hz at each channel. Frequencies with max absolute amplitude from the resulting fits were recorded for each channel and scale parameters were then compared across channels within participants. Three out of fifteen participants were excluded from further analysis because their fitting results yielded no peak in both ISI conditions, i.e., three out of fifteen participants failed to show a peak at the beta frequency, while 11 participants showed similar peaks for both ISI conditions. Lastly, one participant showed a peak in the ISI2 condition but failed in the ISI1 condition. Given the above, the FOI for algorithm detection was centred (± 3 Hz) at the frequency with the channel which contained the largest fitting scale in the ISI2 condition. In addition, this channel served as region of interest (ROI) in the later calculation for detection of beta bursts (Fig. 1E, the left topography showed the ROI for the example; the right topography showed the ROI for all participants, overlapping ROIs were represented by single dot; one ROI was in electrode A2, two ROIs were in D18, three ROIs were in D17, D19, D28, respectively; labels were defined from the 128 channel Biosemi layout, see https://www.biosemi.com/pics/cap_128_layout_medium.jpg).

##### Algorithm for beta burst detection

The detection of beta bursts was based on an established algorithm for detecting fast and slow spindles (Molle et al., 2002; Mölle et al., 2011; Staresina et al., 2015). The algorithm was applied to the trial segmented EEG data starting 5 s before and ending 70 s after the onset of the first tone of a train (see Fig. 1A for an example of raw data). First, trials were filtered with a bandpass filter (FIR filter) of ± 3 Hz centred at the FOI (18.64 Hz in this example, mean FOI is 19.36 Hz) calculated in the above section (Fig. 1B). To prepare the data for burst detection, the root mean square (RMS) of the filtered signal was calculated (Fig. 1C) and smoothed with a moving average of 200 ms. The threshold for burst detection was data driven by finding the maximum correlation between beta burst rate (counts of beta burst) and baseline corrected beta power locked at taps. The correlations were acquired at steps of .25 standard deviations (SD) from 0 to 5 SD (above median), at single trial level and then averaged across participants (Fig. 1D). The resulted maximum correlation was 1.75 SD above the median of filtered signal envelope and was set as the threshold. Events were defined as time points where the resulting RMS signal remained above threshold for 0.1 to 0.5 seconds. For each event, maxima and minima were calculated as peaks and troughs, while the highest peak was designated as the beta burst time (max time). An example of detected beta burst events is demonstrated in Fig. 2.

**Fig. 2.**
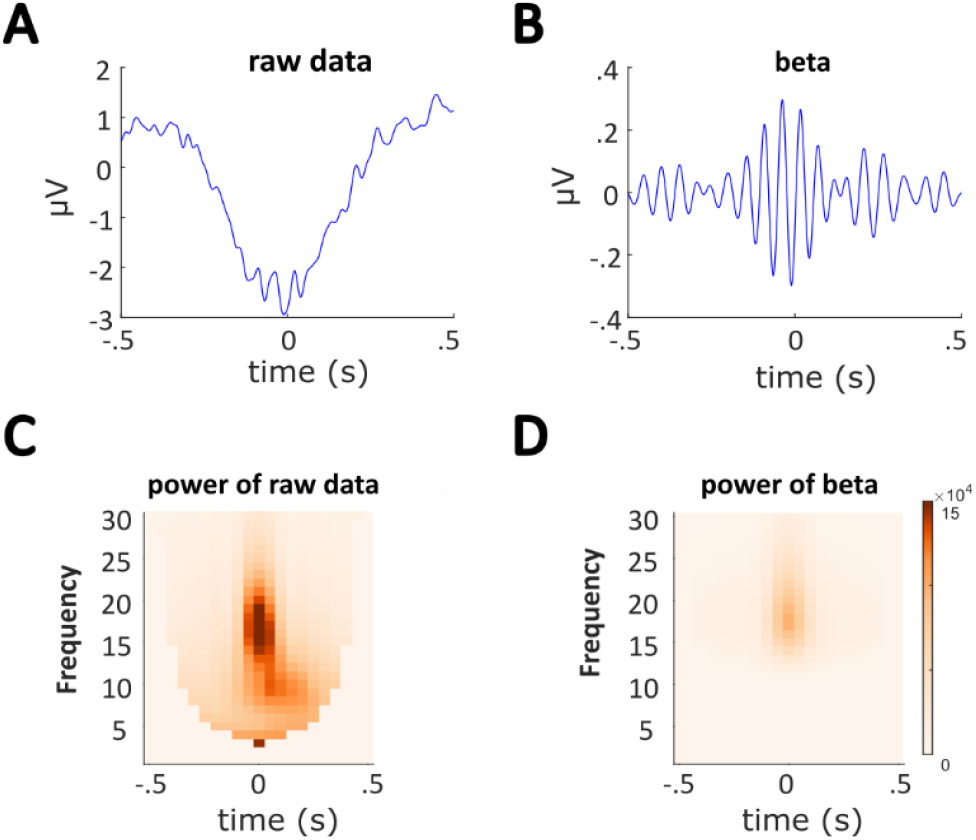
An example of detected beta burst events. Data was incorporating ISI1 and ISI2 conditions and centred at beta burst max time. **(A)** ERPs of Raw data of the detected beta burst events for one subject. Time zero indicates beta burst max time. **(B)** F R bandpass filtered raw data of ± 3 Hz centred at individual frequency of interest (18.64 Hz). **(C)** Time frequency representation of power of raw data (A) from 1-30 Hz revealed a peak at beta range. **(D)** Time frequency representation of FIR filtered data (B).

#### 2.3.4 Comparing Beta Burst Timing Locked to Tap vs. Tone

To explore the relationship between the timing of beta bursts, taps and tones, the timing of beta burst events relative to the tap or tone was calculated separately. First, a time stamp was recorded for the last detected beta burst event that fell within the 1-second window (for ISI1 and 2 seconds for ISI2) before a tap, or a tone, respectively. In other words, when two bursts were detected within the time window before a tap (or a tone), only the last one was counted. A simulated example is illustrated in Fig. 3, which shows time stamps of beta bursts relative to the taps and tone onset time at ISI1. This example illustrates the distribution of beta bursts when they are more locked to the tap, as opposed to the tone time. Next, the standard deviation of the time information aligned to tap/tone was statistically compared between locking to the tap and tone condition using a paired samples t-test. Standard deviation was used as a measure of the distribution of beta burst activity in relation to tap and tone onset, with a lower standard deviation indicating a stronger relationship between beta burst timing to either taps or tones. Additionally, we also carried out a similar analysis taking all beta bursts within 1 second (for ISI1 and two second for ISI2) before a tap, or a tone.

**Fig. 3.**
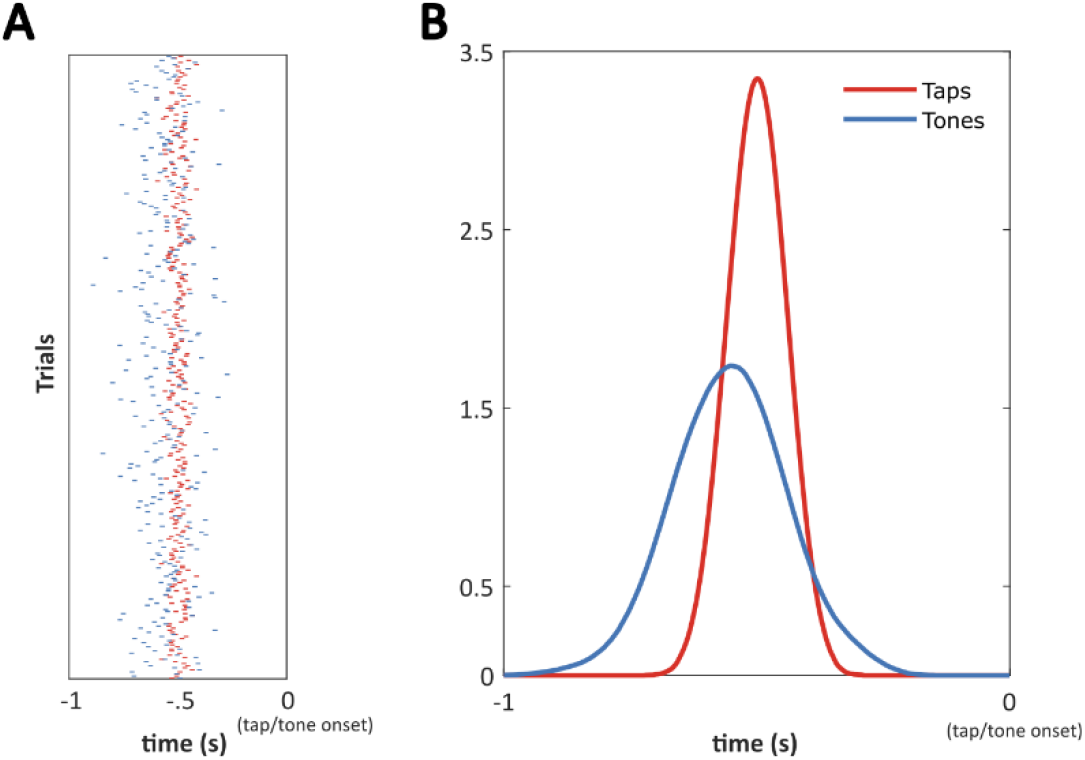
Simulated beta burst events relative to taps/tones time in ISI1. Time stamps of detected beta burst events (max time) locked to taps/tones time were recorded for each subject. **(A)** Distribution of beta burst time stamps across trials locked to taps (red) and tones (blue) time. The example illustrates a case where beta bursts are more closely locked to the taps, than tones. **(B)** Density plot resulting from convolution between Beta burst counts and a gaussian window for taps/tones onset, illustrating higher variance for the tones compared to taps.

#### 2.3.5 ITPC analysis for auditory tracking

After the confirmation that beta bursts track movements (see results 3.2), we aimed to measure neural response tracking of the auditory stimulus. For this purpose, we compared the tone-tap difference in ITPC (inter-trial phase coherence, a measure of the uniformity of phase angles across trials) between the first and last tap/tone trials. During the first few tap/tone trials, as participants were unaware of the stimulation frequency and were still learning the rhythm, the timing of taps varied greatly, although it rapidly improved within three tap/tone trials on average (see Fig. 4B). As revealed by previous findings that phase at low frequencies tracks auditory entrainment (Calderone et al., 2014; Lakatos et al., 2008, 2013; Morillon & Baillet, 2017; Stefanics et al., 2010), phase locking to the tones is predicted to be stronger than that to the taps in the first few tap/tone trials. However, since the taps in the last few tap/tone trials were well timed to the tones, a similar amount of phase locking should be obtained for taps and tones as they essentially became the same event. Hence, a comparison of the tone minus tap ITPC difference (tone-tap ITPC difference) between the first and last five tap/tone trials was taken to reflect auditory tracking. It was expected that the tone-tap ITPC difference would be greater in the first five tap/tone trials than that in the last five tap/tone trials. By contrasting the tone-tap ITPC difference between the first and last five tap/tone trials, this measure theoretically reflects the effect of auditory tracking for the first five tap/tone trials. The number of trials selected was a balance between a quick behavioural adaption found in SMS (see Fig. 4B) and the number of trials needed to establish tracking.

**Fig. 4.**
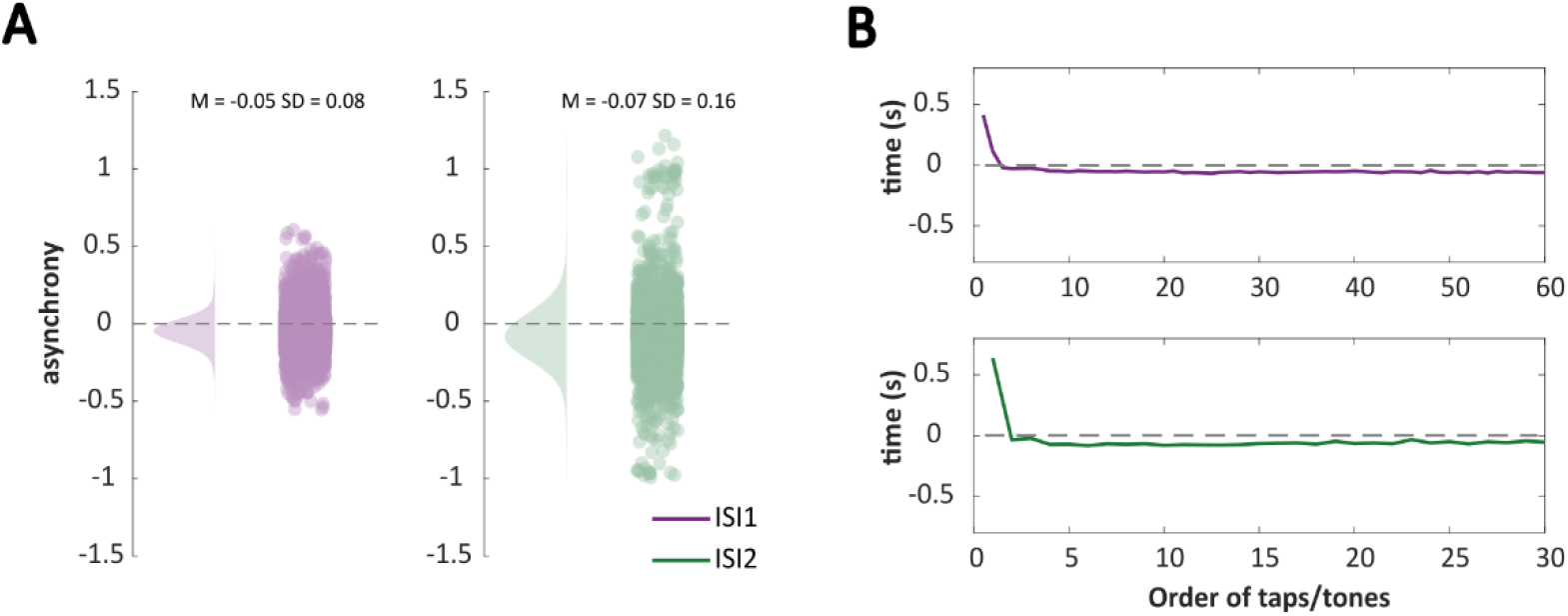
Behavioural asynchrony. The asynchrony performance was calculated by tap minus tone time. Asynchrony of 0 as a baseline was denoted by a dash line. **(A)** Both ISI1 and ISI2 observed a negative mean asynchrony across all subjects (M = −.0459, SD = .0793 for ISI1; and M = −.0693, SD = .1646 for ISI2). Each dot represents one difference between a tone and a tap. **(B)** Time course of asynchrony between tap and tone averaged across trials for ISI1 (60 taps/tones, upper panel) and ISI2 (30 taps/tones, lower panel). The first two taps/tones in the train showed a positive mean asynchrony, but it rapidly declined and stabilized around a negative value for both ISI1 and ISI2.

Preprocessed EEG data was downsampled to 250 Hz and epoched to tap/tone time 10 s before and after onset for the ISI1/ISI2 condition, respectively. Only trials with taps were included in this analysis. In other words, tap/tone trials where subjects missed the tone were excluded. ITPC values for the four conditions (first tones, first taps, last tones, and last taps) were computed by the following procedure individually for ISI1 and ISI2. Complex Fourier-spectra implementing a three-cycle Morlet wavelet time frequency transformation was computed with a moving window of 100 ms (FOI was 0.2 to 1.6 Hz in steps of 0.2 Hz for ISI1 and 0.1 to 1 Hz in steps of 0.1 Hz for ISI2). Next, the resulting complex Fourier-spectra were divided by their amplitude for the calculation of phase angles, which was then summed across trials. This sum of phase angles was then divided by the number of trials to generate ITPC values for each channel. The following analysis used a time window of 2 s centred at tap/tone onset at ISI1 (FOI 1 Hz) and 4 s in ISI2 (FOI 0.5 Hz). For each participant, ITPC locked to first five taps was subtracted from those locked to the first five tones, with this process repeated for the last five taps and tones, resulting in tone-tap differences for the first and last tap/tone. To measure auditory tracking, the difference between the first and last differences at group level was assessed by a one tailed (alpha = 0.01) paired samples cluster-based permutation test (Maris & Oostenveld, 2007) with 2000 randomizations across the time window preselected for ISI1 and ISI2. The significance probability was estimated by the Monte Carlo method.

Phase locking to rhythmic sensory events could be due to entrainment, i.e. locking of a neurophysiological oscillation to an external rhythmic stimulus (Meyer et al., 2020), or it could be due to stimulation artefacts, neural resonance (Helfrich et al., 2019), or superposition of transient event-related potentials (ERPs, Capilla et al., 2011). We explored this issue by measuring entrainment echoes, where phase locking outlasts for several cycles after the termination of a rhythmic stimulus (Hanslmayr et al., 2019). As each trial begins with a 15-second break, this allows enough time for the measurement of echoes. ITPC was calculated 15 seconds before and after the last tone onset across trials for each participant. A three-cycle Morlet wavelet was used to extract phase information from 0.1 to 2 Hz, in steps of 0.1 Hz, with 100 ms time resolution. If entrainment occurs, we expected to observe stronger post-stimulus ITPC at the stimulation frequency comparing to other frequencies. In other words, in ISI1, ITPC at 1 Hz would be stronger than that at 0.5 Hz and vice versa for ISI2. Therefore, a two-way repeated measures ANOVA was carried out to explore the interaction effect between the stimulation frequency conditions (ISI1/ISI2) and the tested frequency (1 Hz/0.5 Hz). A significant interaction effect would be supportive of entrainment. To avoid interference from temporal smearing of the filters applied and the offset effect from the last tone, the time window used for statistical analysis was one to four seconds after the last tone onset for ISI1 and one to seven seconds for ISI2. The selection of time windows of interest allows an observation of three cycles at the FOI.

#### 2.3.6 Calculation of PPC aligned to motor beta burst

Following the identification of neural responses to taps (motor system) and tones (auditory system, see results), it is critical to examine whether there is an interaction between the two systems. To investigate the relationship between motor beta burst timing (max time) and the stimulation frequency (1/0.5 Hz), pairwise phase consistency (PPC) locked to beta burst timing was measured. PPC was preferred over ITPC because it is bias-free with respect to the number of trials and thus allows comparisons between conditions with different numbers of observations (i.e. beta bursts in our case)(Vinck et al., 2010). Preprocessed EEG data with a time window of 14 s, centred at beta burst time for both ISI conditions, were prepared for the computation of PPC. Only the last beta bursts locked to a tap or a tone within 1 second for ISI1 and 2 seconds for ISI2 were included. Phase information was extracted by a three-cycle Morlet wavelet time frequency transformation with a moving window of 100 ms and frequency resolution of 0.1 Hz from 0.1 Hz to 2 Hz and PPC was calculated following the procedure from Vinck et al. (2010). Channel ROIs were selected as the five channels of maximum PPC value averaged across ISI conditions, centred at the beta burst in a time window of 1 second for ISI1 and 2 second for ISI2. Thereafter, a repeated measures ANOVA analysis was carried out to investigate the interaction effect of the ISI condition and the frequency measured (at 1 Hz/0.5 Hz) on PPC. It was expected that the PPC was not only enhanced universally at low frequencies but also affected by the tone frequency (i.e., 1 Hz vs 0.5 Hz), by showing a significant interaction where PPC at the measured frequencies is higher when the measured frequency matches the ISI frequency.

#### 2.3.7 Measuring PPC contrast between high vs. low variance conditions

As the analysis above indicated an interaction between motor and auditory systems (refer to results 3.4), our interest moved to whether such interaction accounts for behavioural differences. Although in finger-tapping tasks, participants often tap before the tone onset, where the time gap is termed NMA (negative mean asynchrony), this gap narrows with training (Repp, 2005; Repp & Su, 2013). Critically, NMA only provides information at the average level, while a large portion of trials (17.67% for ISI1 and 23.25% ISI2) reveals positive asynchrony in our data. This, importantly, could still be interpreted as anticipatory response as it is shorter than the fastest reaction time (Repp, 2005; Repp & Su, 2013). Therefore, to account for the positive asynchrony, we used the absolute value of the asynchrony as a measure of behavioural variance. Tap/tone trials were divided into low and high variance conditions depending on whether the absolute asynchrony was below or above the median for each participant. The exact scheme of PPC calculation from the above section was applied to the low and high variance conditions, respectively. PPC at the stimulation frequency depending on ISI condition (1 Hz and 0.5 Hz was measured for ISI1 and ISI2, respectively) were statistically compared between low and high asynchrony conditions with repeated measures ANOVA.

### 2.4. Data and code availability

The pre-processed EEG data and the code used to generate the results reported in this manuscript can be downloaded from https://osf.io/ancqm/?view_only=3099faaab84e4713828991a104801259

## 3. Results

### 3.1 Behavioural results

Three out of fifteen participants were excluded from further analysis because their fitting results yielded no peak in both ISI conditions at the preparation of beta burst detection, i.e., three out of fifteen participants failed to show a peak at the beta frequency. This left 12 participants in the following analysis. The asynchrony calculated by tone minus tap time revealed a negative mean asynchrony (Fig. 4A) for both ISI1 (M = −.0459, SD = .0793, N = 13068, unit in second) and ISI2 (M = −.0693, SD = .1646, N = 6513). There was less asynchrony variance in ISI1 compared with ISI2. In 17.67% and 23.25% of tap/tone trials we observed positive asynchrony in ISI1 and ISI2, respectively. The asynchrony declined quickly within trials (i.e., a series of 60/30 taps for ISI1 and ISI2 respectively), indicating rapid adaptation to the entraining frequency, despite participants being unaware of the ISI condition. Noticeably, this rapid adaptation occurs within only three taps/tone iterations on average (see Fig. 4B).

### 3.2 Beta bursts are stronger locked to taps than tones

After EEG preprocessing, a mean number of 39 (from 24 to 56) trials remained for later analysis. ISI1 and ISI2 contained a same average number of trials (19, both ranging from 11 to 29). There was an average of 263 (from 68 to 376) beta burst events locked to tones detected in ISI1 and 268 (from 133 to 479) events in ISI2. For beta burst events locked to taps, this number is 263 (from 66 to 377) in ISI1 and 268 (from 135 to 478) in ISI2. There is no difference regarding the number of beta bursts between trials locked to taps and tones (*t* (11) = −.3427, *p* = .7383 for ISI1 and *t* (11) = −.4419, *p* = .6671 for ISI2). To investigate the timing relationship between beta bursts and taps or tones, the distribution of beta burst timing locked to taps and tones was compared by examining the standard deviation of the beta burst time stamps. A higher standard deviation was interpreted to reflect greater temporal variance and therefore less locking to the taps or tones. Findings indicated that the standard deviation was lower when beta bursts were locked to taps as compared with tones at both ISI1 and ISI2 (*t* (11) = − 2.0266, *p* = .0676 for ISI1 and *t* (11) = −3.2607, *p* = .0076 for ISI2). Despite the marginal significance in ISI1, the effect was further confirmed by a repeated measures ANOVA, indicating that beta burst timing tracked the taps significantly better than tones in both ISI conditions (*F* (1, 11) = 10.1313, *p* =.0087), see Fig. 5). Surprisingly, we also observed a main effect of ISI condition, where ISI1 demonstrated a closer relationship between burst, tap and tone timing (*F* (1,11) = 210.5132, *p* =.0000). Although we were unable to disentangle the relationship of beta burst timing and taps from potential interference from tones, the observed pattern across ISI conditions suggests that beta burst timing is more closely related to the movement than to the periodic auditory stimuli.

**Fig. 5.**
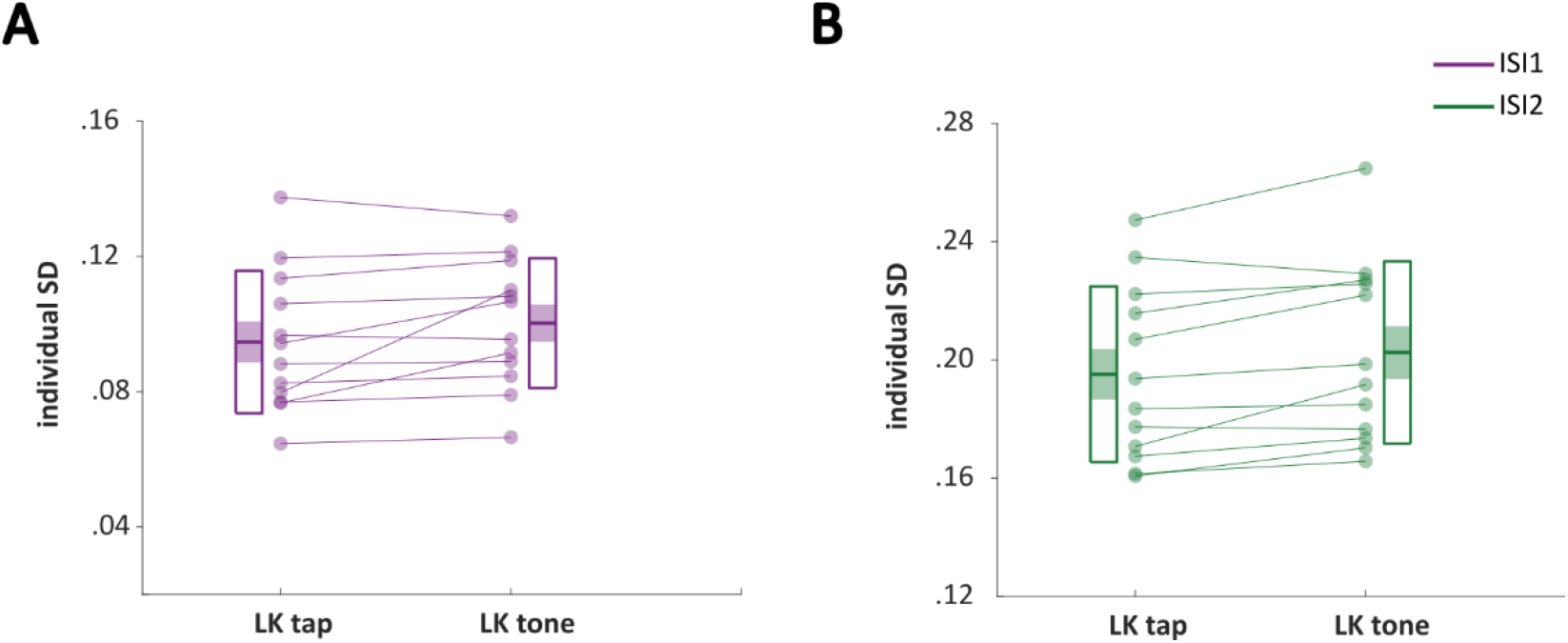
Comparisons of distributions of beta burst events between tap and tone onset. Standard deviation of beta burst time stamps locked to tone/tap were calculated for each subject for ISI1 **(A)** and ISI2 **(B)**, respectively. Each dot represents one subject. Within subject tendency between tap and tone was showed by linked dots. The top and bottom line of the boxes represent 95% percentiles above and below the mean (middle line of the boxes). Shaded areas denote SEM. A paired samples t-test confirmed that the beta burst timing tracks taps significantly better than tones in both ISI conditions. Although the difference was marginally significant in ISI1, the main effect of burst timing locked to tap vs. tone was confirmed by a repeated measures ANOVA taking both ISI conditions

Additional analysis counting all beta bursts before the same time window before a tap (or a tone) revealed similar results but with weaker effects (*t* (11) = −1.6374, *p* = .1298 for ISI1 and *t* (11) = − 2.3839, *p* = .03266 for ISI2).

### 3.3 Phase of stimulation frequency is more strongly locked to tones than to taps, but no evidence for entrainment

After confirming a relation between beta burst and tap timing we aimed to measure the neural correlates for auditory tracking. Participants were not informed of the stimulation frequency before the start of each trial. Therefore, the timing of taps in the first few tap/tone trials varied greatly. Given the findings that the ongoing phase tracks auditory stimuli (Calderone et al., 2014; Lakatos et al., 2008, 2013; Morillon & Baillet, 2017; Stefanics et al., 2010), ITPC locked to tones was expected to be stronger compared to ITPC locked to taps particularly in the first few tap/tone trials. In contrast, ITPC for taps and tones should not differ in the last few tap/tone trials, because of the nearly perfect timing of taps towards the end of the trial (see Fig. 4B). Hence, the auditory tracking could be measured by comparing the tone-tap ITPC difference between the first and last five tap/tone trials. We used five trials for this comparison because it represented a good balance between a quick behavioural adaption found in SMS (see Fig. 4B) whilst allowing for long enough segments to establish tracking. Following this logic, auditory tracking was evaluated by a cluster-based paired-samples permutation test in the comparison of tone-tap ITPC difference between the first and last five tap/tone trials. There was an average of 74 (ranging from 44 to 109) and 79 (ranging from 52 to 105) tap/tone trials involved in the calculation of ITPC locked to the first 5 tap/tone in ISI1 and ISI2, respectively. In the last five tap/tone condition, there were 91 (ranging from 32 to 145) and 91 (ranging from 54 to 135) tap/tone trials available for calculating ITPC. This analysis resulted in two clusters (channels highlighted in Fig. 6A and B) with significant t-values (*p_corrected_* = .0100 in ISI1 and *p_corrected_*= .0014 in ISI2), indicating stronger phase-locking to the tone at the beginning of a trial compared to the end of the trial. As predicted, the averaged ITPC within clusters for the first tones is higher than that for the first taps, while the last tones and last taps showed similar ITPC (Fig. 6C and D for ISI1 and ISI2, respectively). Together, these results show that phase tracks the auditory activity elicited by the tones, as opposed to motor activity elicited by taps.

**Fig. 6.**
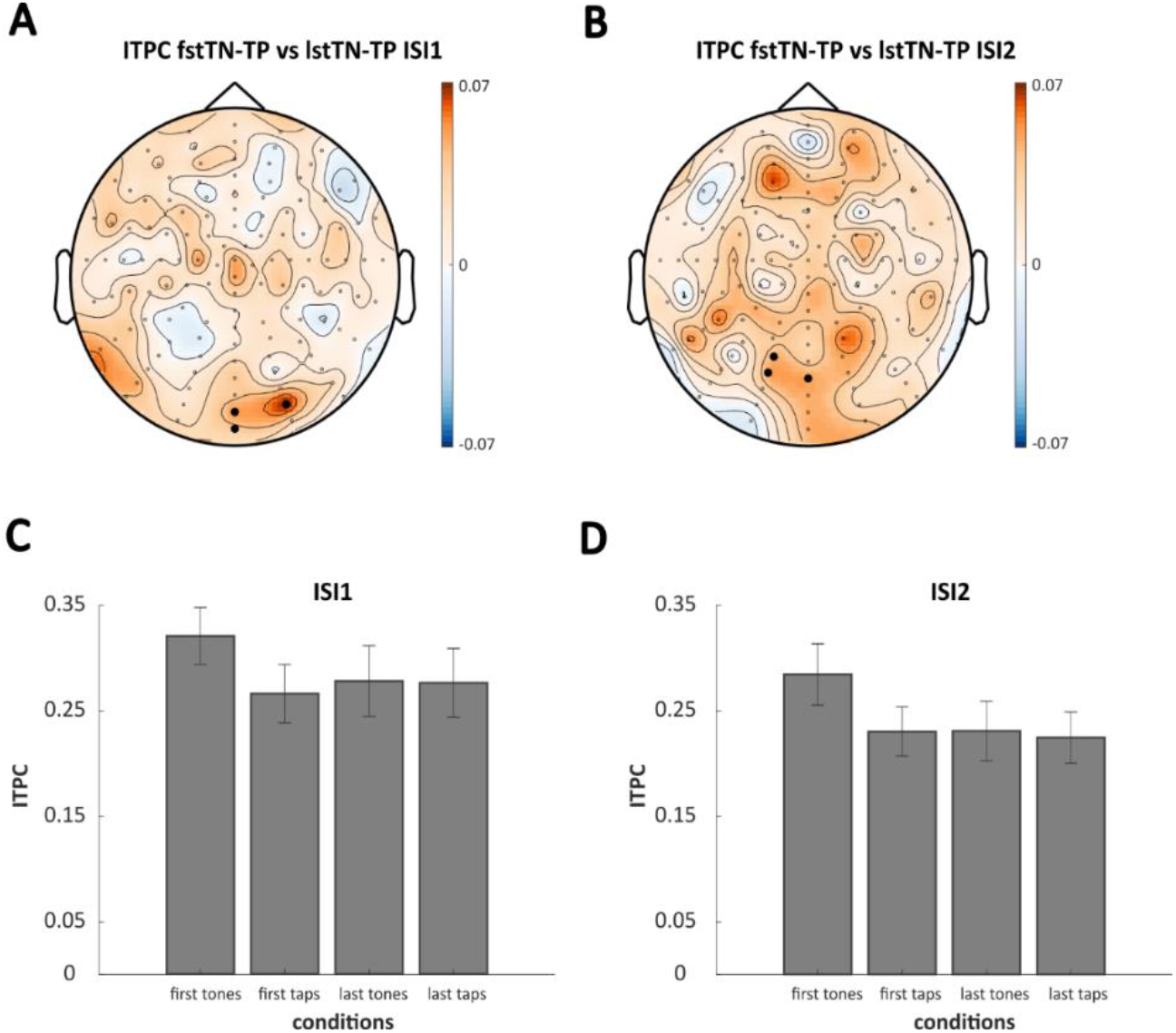
Auditory tracking. The auditory tracking of tones was measured by the comparisons of the tone minus tap ITPC difference (tone-tap ITPC difference) between the first and last five taps/tones of a train. **(A)** and **(B)** Cluster-based permutation test indicated statistically significant differences between the first and last five taps/tones of a train regarding tone-tap ITPC difference. Electrodes with the most pronounced differences were highlighted. **(C)** and **(D)** ITPC averaged across highlighted electrodes in (A) for ISI1 and (B) for ISI2 were calculated for the first five taps/tones, respectively. In both ISI1 and ISI2, the averaged ITPC for the first tones is the highest comparing with first taps, last tones, and last taps.

To establish whether the phase locking to the auditory stimulation frequency was due to neural entrainment, we measured entrainment echoes, i.e., where phase locking lasts for several cycles even after the rhythmic stimulus stops (Albouy et al., 2017; Hanslmayr et al., 2014). ITPC from 0.2 to 2 Hz is shown before and after the last tap onset (time 0) in Fig. 7A and B. Support for entrainment would be if post-stimulus ITPC is stronger at the stimulation frequency compared to other frequencies. Hence, post-stimulus ITPC in ISI1 and ISI2 at the two tested frequencies (0.5 and 1 Hz) were compared using a two-way repeated measures ANOVA. Statistical results revealed a significant main effect of ISI condition (*F* (1,11) = 8.2960, *p* = .0150) such that ITPC is stronger in ISI1 compared with ISI2 for both tested frequencies. This was further confirmed by Bayes factor indicating evidence for H_+_; specifically, BF_+0_ = 11.52, which means that the data are approximately 11.52 times more likely to occur under H_+_ than under H_0_. The error percentage is 1.207%, which indicates a good stability in obtaining the result. However, the non-significant interaction between ISI condition and tested frequency (*F* (1,11) = 2. .8224, *p* = 0.1211, BF_+0_ = .878) does not lend support to an entrainment process).

**Fig. 7.**
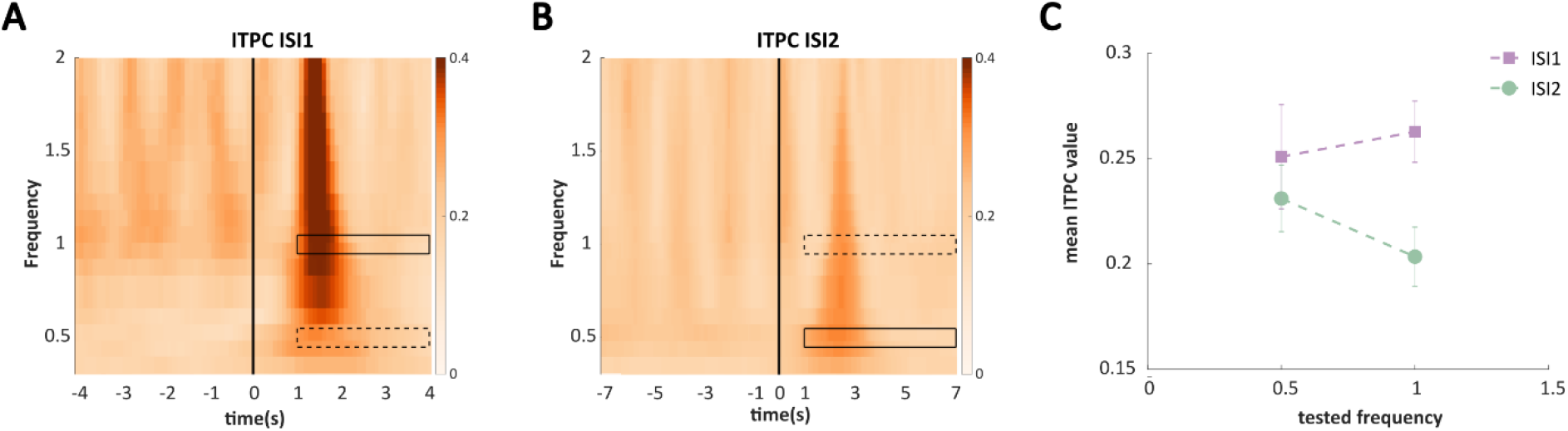
ITPC before and after the last tone. **(A)** ITPC from 0.2 to 2 Hz in ISI1, in a time window of four seconds before and after the last tone onset (time 0, as indicated by the solid vertical line). **(B)** Same as (A) but for ISI2 and with a time window of seven seconds before and after the last tone onset. Black boxes in both (A) and (B) indicate TOI and FOI used for the two-way repeated measures ANOVA. While the solid-line boxes indicate congruence between stimulating and tested frequency, the dashed-line boxes indicate incongruence between them. **(C)** Averaged ITPC within corresponding FOI and TOI in both stimulating and tested frequencies. The two-way repeated measures ANOVA revealed a significant main effect of the stimulation frequency conditions, showing stronger ITPC in ISI1 comparing to ISI2. No significant interaction as evidence of entrainment was observed.

### 3.4 Beta bursts are locked to the phase of the auditory stimulation frequency

The above analyses suggest that the timing of beta bursts tracked tapping, whereas the phase at the stimulation frequency tracked the auditory stimulus. Since the task required participants to synchronise their taps to the stimulation frequency, we analysed if the timing of beta bursts is locked to the phase of the auditory stimulation frequency. Such phase locking would be an indication for an interaction between the motor (movement, i.e., taps) and sensory (auditory, i.e., tones) systems. To this end, we calculated the pair-wise phase consistency (PPC, a measure of phase locking that is unaffected by different trial numbers; Vinck et al., 2010), averaged across all channels, time-locked to beta bursts from 0.1 to 2 Hz. ROIs for statistical analysis were then later chosen as the five channels with maximum PPC values averaged across ISI conditions (Fig. 8A). This analysis revealed a peak at the stimulation frequency for both ISI1 and ISI2 conditions (Fig. 8B). This finding was further confirmed by the result of a two-way ANOVA revealing a significant interaction between ISI condition (ISI1 vs. ISI2) and PPC at the measured frequency (1 Hz vs. 0.5 Hz; *F* (1,11) = 8.1009, *p* =.0159), suggesting that beta burst phases were specifically locked to the stimulation frequency. No significant main effects were found. Grand averaged PPC across participants show peak PPC at central areas (Fig. 8A) for both ISI1 and ISI2. Together, these results show that beta bursts are phase locked to the stimulation frequency, suggesting a synchronization between the motor and the auditory systems during SMS.

**Fig. 8.**
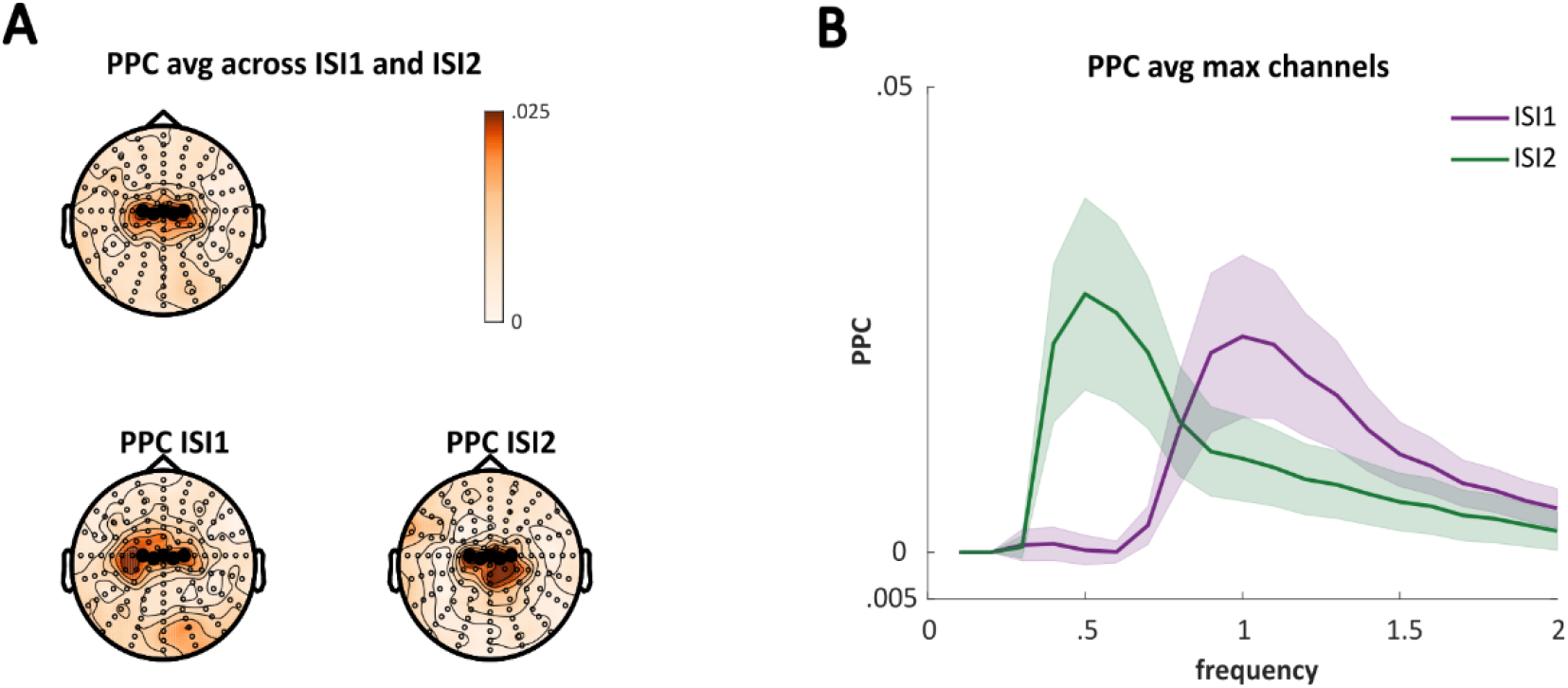
PPC locked to motor beta burst. **(A)** Topographical distribution of the grand averaged PPC showing peaks at central areas in both ISI conditions (lower panels) and the average across conditions (top panel). The five maximum channels averaged across ISI conditions were used as ROI for statistical analysis (highlighted). **(B)** Pairwise phase consistency (PPC) time-locked to beta burst averaged at the ROI from 0.1 to 2 Hz is shown. Peak PPC can be observed at stimulation frequency in both ISI1 (purple line, peaks at around 1 Hz; shaded areas indicate SEM) and ISI2 (green line, peaks at around 0.5 Hz).

### 3.5 Timing of beta bursts explains variance in behavioural performance

The above results showing beta burst phase locking to the stimulation frequency, point towards an interaction between the neural response accounting for movements and the auditory stimulus. If this interaction indeed reflects synchronization between the motor and auditory system, then this effect is expected to correlate with variations in tapping behaviour. To test this, tap/tone trials were conditioned as low and high variance taps (i.e., good or bad timing, respectively), depending on whether the absolute value of asynchrony was above or below the median across trials (absolute asynchrony) for each participant. PPC locked at beta burst was then compared between low and high variance trials at the stimulation frequency of ISI condition using a repeated measures ANOVA. As expected, this analysis revealed stronger PPC in the low compared to high variance condition (*F* (1,11) = 5.2966, *p* =.0419, Fig. 9), indicating that beta bursts tend to peak at a consistent phase at the stimulation frequency when taps are temporally more accurate. Surprisingly, we also observed a main effect of ISI condition (*F* (1,11) = 4.9190, *p* =.0485), but no interaction.

**Fig. 9.**
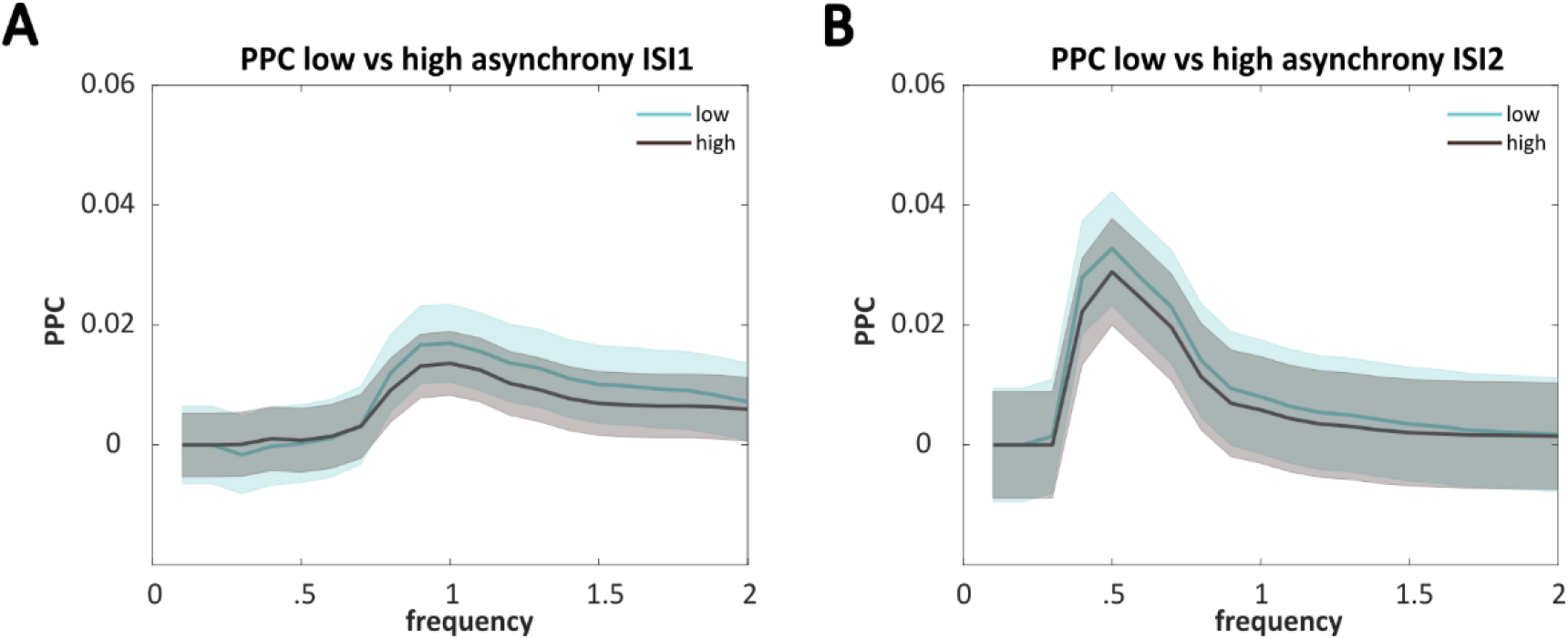
Comparisons of PPC between low and high asynchrony conditions. PPC time-locked to motor beta burst was compared between low and high asynchrony conditions, which was conditioned by individual median of tap-to-tone absolute asynchrony. Shaded areas indicate SEM. **(A)** The difference in PPC from 0.1 to 2 Hz peaked at the entrainment frequency in ISI1 (peaked at 1 Hz) and **(B)** in ISI2 (peaked at 0.5 Hz).

Together, these results indicate that beta burst timing is closely related to taps and is phase locked to the stimulation frequency of the auditory stimulus. Importantly, beta burst phase locking accounts for tapping variations on a trial-by-trial level. These results suggest that phase-locked beta bursts may act as an internal link between the motor and auditory systems during SMS.

## 4. Discussion

Accumulating evidence suggests that both rhythmic sensory stimulation and the timing of movements contribute to SMS, which requires interaction between the motor and auditory systems. However, the underlying neural correlates remain unclear. We hypothesized that the timing of beta bursts and the phase of the stimulation frequency play an important role in the synchronization of the motor and auditory systems and that an interaction between beta burst timing and the phase of the stimulation frequency relates to behavioural performance. To test our hypothesis, we explored the neural representation for taps and tones, which were related to beta burst timing and auditory tracking, respectively. Importantly, we observed an interaction between beta burst timing and phase locking at the stimulation frequency (1 Hz for ISI1 and 0.5 Hz for ISI2), where the degree of such phase locking correlated with behavioural performance. Together these results suggest a functional role of beta bursts and their phase locking at the stimulation frequency for synchronizing movements to a rhythmic sensory stimulus.

Behaviourally, the finding of negative mean asynchrony (NMA) is consistent with previous findings (Repp, 2005; Repp & Su, 2013). In general, participants adapted to the rhythm quickly, and anticipated the next tone within three stimuli. Additionally, tapping performance is generally better in ISI1 compared to ISI2 by showing a shorter NMA and less variance in tap-tone gaps. These behavioural results indicate higher difficulty in ISI2, consistent with previous findings showing that when ISI is longer than 1800 ms, performance drops significantly (Miyake et al., 2004). Although approximately 20% of trials showed a positive mean asynchrony, i.e., 20% taps occurred following tone onset, reaction times shorter than 150 ms (about the shortest reaction time, Repp, 2005; Repp & Su, 2013) should still be considered as predictive behaviour as opposed to reactive response. When this is taken into consideration, the proportion of trials with taps following tones by more than 150 ms is similar between ISI1 (5.85%) and ISI2 (6.26%).

Before studying the interaction between motor and sensory systems during SMS, it is critical to investigate the mechanisms underlying the phase locking to the auditory stimuli. Measuring entrainment echoes, where phase locking continues for several cycles after the termination of a rhythmic stimulus is one way of disentangling neural entrainment from other phase-locked neural responses (Hanslmayr et al., 2019). However, our results show no evidence for entrainment. ITPC in ISI1 after stimulus offset was higher than ISI2 in general, indicating that the stimulation frequency of .5 Hz may be too slow to trigger strong entrainment. Interestingly, the ITPC at 1 Hz is significantly higher in ISI1 comparing to ISI2 (*t* (11) = 3.4008, *p* = .0059), but not at 0.5 Hz. Future designs may employ higher frequencies to test entrainment. However, it is also a challenge to distinguish whether the functional role of beta bursts, i.e., a prediction role as part of pre-movement activity, or other functional roles (e.g., facilitating responses back to baseline) as post-movement response. This becomes more of a concern when the stimulation rate increases.

Beta oscillations (13-30Hz) are considered a default mode of the motor system (Niso et al., 2016) and have been shown to be related to the representation of temporal information, with (Arnal et al., 2015; Arnal & Giraud, 2012; Kulashekhar et al., 2016; Morillon & Baillet, 2017) and without overt movements (Fujioka et al., 2012, 2015; Morillon & Baillet, 2017). Classically, beta activity is characterized by power decreases or event-related desynchronization (ERD) before movement, and power increases (event-related synchronization, ERS) gradually back to or even above baseline following the movement (van Wijk et al., 2012; Pfurtscheller & Da Silva, 1999). However, the conventional method of averaging beta signals across trials was recently challenged by studies investigating trial-wise dynamics of beta oscillations (Feingold et al., 2015; Little et al., 2019; Shin et al., 2017). These studies suggest that discrete, high amplitude, transient (typically last <150 ms, Feingold et al., 2015) and infrequent beta bursts, which are tightly locked to movements, are dominant in motor systems across humans and animals (Sherman et al., 2016). Importantly, transient beta fluctuations, which could be masked out in conventional power observation, can help the discrimination of brain states associated within movements (West et al., 2022). This research motivated us to investigate the relationship between beta burst timing with tap and tone timing. In parallel with the better behavioural performance in ISI1, beta burst timing was more closely related to the taps (and tones) in ISI1 compared to ISI2 (Fig. 5). One may interpret such a result as beta burst timing predicting better behavioural performance. However, on the one hand, since taps were also closely tracing tones, we were unable to disentangle the effects of tones and produce a pure effect of beta burst timing on taps. On the other hand, while the shorter averaged gap between burst timing before tap in ISI1 than ISI2 relates to better performance, this is in violation with earlier findings showing that earlier pre-movement beta timing behaviourally correlates with fast responses (Little et al., 2019). To reconcile this discrepancy, here our results suggest that in SMS, it is the interaction between the sensory tracking phase and beta burst timing correlates with prediction performance, rather than beta burst timing per se.

By analysing directed functional connectivity, previous studies found that even during passive listening with no movements, left sensorimotor (LSM) beta (18-24 Hz) directs towards the right associative cortex (rAA) in a temporal prediction task. Reversely, neural responses corresponding to auditory stimulus rates directs from rAA to LSM (Morillon & Baillet, 2017). However, whether such directional interplays relate to the prediction of behavioural performance remains unknown. Despite the equivocal role of beta burst timing in prediction of tapping performance, we found that the beta burst locking to the stimulation phase was functionally correlated with better behaviour in both ISI conditions. Limited by the EEG method we used in this study, we were unable to investigate the origin of beta bursts and the time course of interactions between motor and sensory cortices during SMS. Noticeably, in our study with movements, the frequency band of beta ranges from 18-23 Hz (except one subject at 13 Hz), similar to the beta range revealed by the above-mentioned study examining functional connectivity with passive listening (Morillon & Baillet, 2017).

Although delta (<4 Hz, Molinaro et al., 2018) entrainment was frequently studied, it was scarcely studied at frequencies less than 1 Hz in auditory entrainment. Our experiment contained two conditions implementing inter stimulus intervals of one and two seconds, corresponding to 1 and .5 Hz, respectively. Although in both ISI conditions we observed a stronger beta burst phase locking at the stimulation frequency related with shorter tap-tone gap, PPC in general is higher for ISI2 compared to ISI1, regardless to tapping accuracy. There is little existing evidence to explain this result. On a behavioural level, prediction performance in finger tapping drops when ISI is greater than 1800 ms compared to the control condition (Miyake et al., 2004). One possible explanation is that the tapping prediction may be more reliant on the phase locking between the beta burst and the stimulus when the task becomes more challenging.

In summary, our study found that the degree of phase-locking of beta burst timing to the auditory stimulation rate accounts for the accuracy of predictive tapping at the single-trial level. These results provide first evidence supporting a functional role of beta burst timing in the coordination between motor and auditory systems during SMS. However, the origin of this functional beta burst and the dynamic interaction between the motor and auditory systems during prediction need further exploration.

## Acknowledgements

We thank Verena Braun for collecting the data. SH was supported by grants from European Research Council (ERC) Consolidator Grant 647954, and the Wolfson Foundation and Royal Society. SH and KS were funded by a grant from the Economic and Social Research Council (ESRC; ES/R010072/1). CM was granted by the Wellcome Trust (ISSF; 1516ISSFIRA4). QC received funding from the China Scholarship Council (CSC).

The authors declare no competing financial interests.

## Notes

https://osf.io/ancqm/?view_only=3099faaab84e4713828991a104801259

### Competing Interest Statement

The authors have declared no competing interest.

